# Connectome and microcircuit models implicate atypical subcortico-cortical interactions in autism pathophysiology

**DOI:** 10.1101/2020.05.08.077289

**Authors:** Bo-yong Park, Seok-Jun Hong, Sofie Valk, Casey Paquola, Oualid Benkarim, Richard A. I. Bethlehem, Adriana Di Martino, Michael P. Milham, Alessandro Gozzi, B. T. Thomas Yeo, Jonathan Smallwood, Boris C. Bernhardt

## Abstract

Both macroscale connectome miswiring and microcircuit anomalies have been suggested to play a role in the pathophysiology of autism. However, an overarching framework that consolidates these macro and microscale perspectives of the condition is lacking. Here, we combined connectome-wide manifold learning and biophysical simulation models to understand associations between global network perturbations and microcircuit dysfunctions in autism. Our analysis established that autism showed significant differences in structural connectome organization relative to neurotypical controls, with strong effects in low-level somatosensory regions and moderate effects in high-level association cortices. Computational models revealed that the degree of macroscale anomalies was related to atypical increases of subcortical inputs into cortical microcircuits, especially in sensory and motor areas. Transcriptomic decoding and developmental gene enrichment analyses provided biological context and pointed to genes expressed in cortical and thalamic areas during childhood and adolescence. Supervised machine learning showed the macroscale perturbations predicted socio-cognitive symptoms and repetitive behaviors. Our analyses provide convergent support that atypical subcortico-cortical interactions may contribute to both microcircuit and macroscale connectome anomalies in autism.

## Introduction

Autism is one of the most common neurodevelopmental conditions, with persistent impairments that often challenge affected individuals, their families, health care, and educational system at large ^1–3^. Despite extensive research efforts, the conceptualization and management of autism continue to face significant challenges. A major difficulty in the neurobiological understanding of autism is that the condition impacts multiple scales of brain organization ^4–10^. Contemporary neuroimaging studies suggested that autism is characterized by both macroscale anomalies in brain connectivity ^6–8,10–14^ and local changes in microcircuit function such as excitation/inhibition imbalance ^4,5,9,15–17^. However, we currently lack an overarching framework that can bridge the topographical changes at macroscale and microscale dysfunctions in autism pathophysiology. Here, we examine how atypical macroscale organization relates to microcircuit imbalances in autism.

Systems neuroscience has recently gained unprecedented opportunities to interrogate the human brain at multiple scales in both health and disease ^4,6,8,9,18^. Although a wealth of studies in autism spectrum conditions have examined local disturbances in cortical morphology ^12,19^ and functional connectivity ^6,10,14,20,21^ as well as mesoscale functional miswiring ^22–25^, less is known about macroscopic changes in structural connectivity ^26–29^. Previous studies observed atypical diffusivity parameters and streamline connections in local brain regions and pathways in autism ^27,28^. Such information can be inferred via diffusion magnetic resonance imaging (dMRI) and tractographic algorithms that reconstruct structural wiring ^30–35^. Macroscale brain organization is increasingly studied using manifold learning techniques that projects high dimensional connectome descriptions into low dimensional representations. In neurotypical individuals, these techniques have gained significant traction to study large-scale principles of functional and microstructural neuroimaging data, owing to its advantage of representing complex pattern of brain connectivity in a compact analytical space ^6,36–39^. This approach, however, remains underexplored in the study of structural connectivity based on dMRI in general, and the assessment of structural wiring perturbations in autism in particular.

A further deliverable of determining principles of structural connectivity is the ability to predict functional dynamics from structural information ^18,40–43^. One class of methods simulates whole-brain functional dynamics via a network of anatomically connected neural masses ^41–43^. In contrast to approaches that assess structure-function coupling statistically ^44–47^, these models are governed by biophysically plausible parameters that are anchored in established models of neural circuit function ^18,40^. A recent study in healthy young adults established that these models robustly simulate intrinsic functional networks from structural connectivity data, and model inversion approaches can be used to estimate regionally varying microcircuit parameters, specifically recurrent excitation/inhibition and external subcortical input into cortical microcircuits ^18^. Thus, applying these novel models to autism will lend additional insights into microcircuit-level correlates of macroscale connectivity perturbations.

Our study aimed to understand the relationship between macro- and microscale perturbations in autism relative to neurotypical controls. We applied manifold learning techniques to dMRI data to generate low dimensional structural connectome representations ^39,48^, and used these to build a macroscale account of topographical structural divergence in autism. Biophysical computational simulations were then used to infer microcircuit-level imbalances at a regional level, specifically recurrent excitation/inhibition and excitatory subcortical input ^18^. We embedded our results in a neurobiological and neurodevelopmental context by decoding the macroscale patterns against post-mortem maps of gene expression data ^49–51^. Finally, we established associations between our macroscale findings and autism symptom severity via using supervised machine learning with five-fold cross-validation.

## Results

Our sample consisted of 47 individuals with autism and 37 neurotypical controls obtained from the two independent sites (see **Table S1** for demographic information) from the Autism Brain Imaging Data Exchange initiative (ABIDE-II; http://fcon_1000.projects.nitrc.org/indi/abide) ^52,53^. See *Methods* for details on participant selection, image processing, manifold generation, computational modeling, transcriptomic decoding, and statistical analysis as well as symptom prediction.

### Large-scale structural connectome manifolds in neurotypicals

We estimated a cortex-wide structural connectome manifold using non-linear manifold learning (https://github.com/MICA-MNI/BrainSpace) ^39^. The template manifold was estimated from an unbiased and group representative structural connectome ^54^ to which individual manifolds were aligned (see *Methods*) ^39^. The three dimensions (henceforth, M1, M2, and M3) reflected the principle axes of variation in structural connectivity with 50.6% variance (**Fig. 1A**), whereby each cortical region is described by its position along these axes. The individual dimensions extended from somatomotor to visual areas (M1), differentiated lateral parietal and prefrontal cortices (M2), and showed a lateral to medial cortical axis (M3; **Fig. 1B**).

**Fig. 1.**
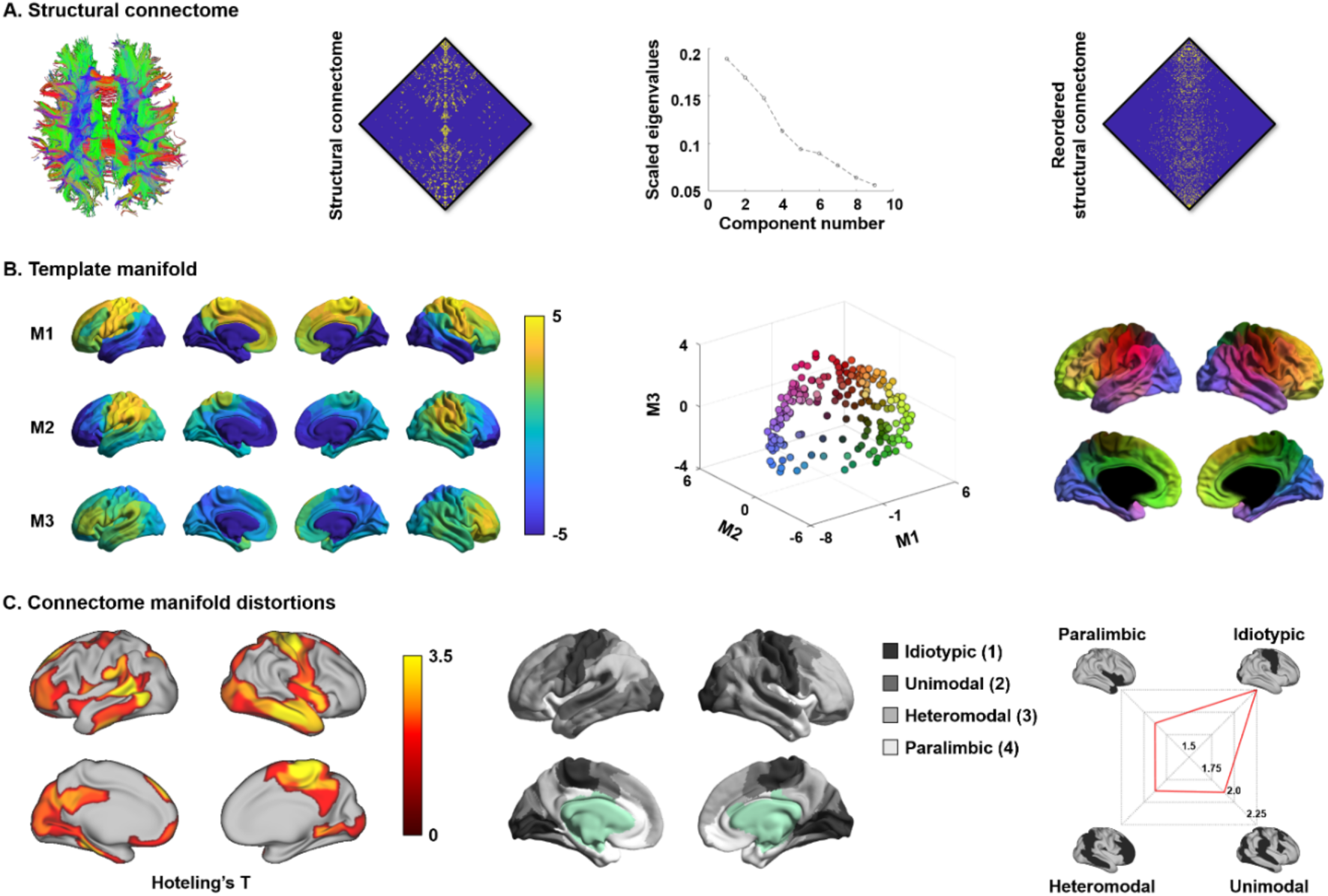
Structural connectome manifolds. **(A)** Fiber tracts generated from dMRI, a cortex-wide structural connectome, and a scree plot describing connectome variance after identifying principal eigenvectors. The structural connectome reordered according to M1 is shown on the right side for better visualization. **(B)** Manifolds estimated from the structural connectome. Three dimensions (M1, M2, and M3) explained >50% of variance and corresponded to the clearest eigengap. Each data point (*i*.*e*., brain region) was represented in the three-dimensional manifold space with different color, and it was mapped onto the brain surface for visualization. **(C)** The t-statistics of the identified regions that showed significant between-group differences in these dimensions between individuals with autism and controls. Findings have been corrected for multiple comparisons at FDR < 0.05. Stratification of between-group differences effects along cortical hierarchical levels (*i*.*e*., middle column) ^55^ is presented in the radar plots. *Abbreviation:* FDR, false discovery rate.

### Connectome manifold distortions in autism

Cortex-wide multivariate analyses compared manifolds spanned by M1–M3 between individuals with autism and controls, using a model that controlling for age, sex, and site in addition to including group effects. We observed macroscale distortions in autism in multiple networks, with primary effects in sensory and somatomotor as well as heteromodal association cortices (false discovery rate (FDR) < 0.05; **Fig. 1C**). Stratifying effects according to a seminal model of neural organization that contains four cortical hierarchy levels (1: idiotypic; 2: unimodal association; 3: heteromodal association; 4: paralimbic) ^55^, we identified peak effects in idiotypic areas followed by unimodal and heteromodal association cortices. Similarly, when analyzing effects with respect to seven intrinsic functional communities ^56^, we also observed strongest between-group differences in somatomotor networks followed by higher-order systems such as the default-mode network (**Fig. S1A**).

Expressing multivariate changes in terms of manifold expansion/contraction ^57^, we found regionally-variable patterns with somatomotor and posterior cingulate cortices showing contractions while heteromodal association cortex underwent expansions in autism relative to controls (**Fig. S1B**). Similar findings were observed in both sites (**Fig. S1C**). In addition, we assessed the degree of head motion of each individual during the dMRI scan based on framewise displacement (FD), and found that the mean FD was not different between autism and controls (p > 0.3) (**Fig. S2A**). Notably, between-group differences in structural manifolds were virtually identical when additionally controlling for mean FD, indicating the head motion did not affect patterns of structural connectome perturbations in autism (**Fig. S2B**). Repeating analyses separately in children (age < 18) and adults (age ≥ 18), effects in adults with autism were highest in higher-order frontoparietal/paralimbic areas while children with autism displayed strongest anomalies in somatomotor/idiotypic regions (**Fig. S3**), similar to age-stratified results in previous functional connectome findings ^6^.

Prior research has suggested atypical cortical morphology in autism, showing anomalies in both cortical thickness and folding relative to controls ^12,19^, motivating an assessment of morphological effects on manifold findings. Correlating manifold distortions with cortical thickness and folding variations, we observed only marginal relations (p > 0.1; **Fig. S4A**). In addition, connectome manifold differences between autism and controls were still measurable when controlling for cortical thickness and curvature in the same model, indicating that structural connectome perturbations occurred above and beyond any potential variations in cortical morphology (**Fig. S4B**).

### Microcircuit parameters from biophysical network modeling

Biophysical computational simulations ^18^ were employed to complement our macroscale findings by modelling atypical microcircuit-level functional dynamics in autism. By linking ensembles of local neural masses (*i*.*e*., theoretical cell population models for excitatory neurons which reciprocally inhibit each other; **Fig 2A**) with diffusion-derived long-range structural connectivity, our computational model simulated dynamic functional time-series, which allowed for the simulation of whole-brain functional connectivity. Notably, by iteratively tuning the parameters of the local neural masses, our simulation generated maximally similar functional connectivity patterns compared to experimental data, which also resulted in an optimal set of biophysical parameters (specifically, recurrent *excitation/inhibition* and excitatory *subcortical/external input;* see *Methods*).

**Fig. 2.**
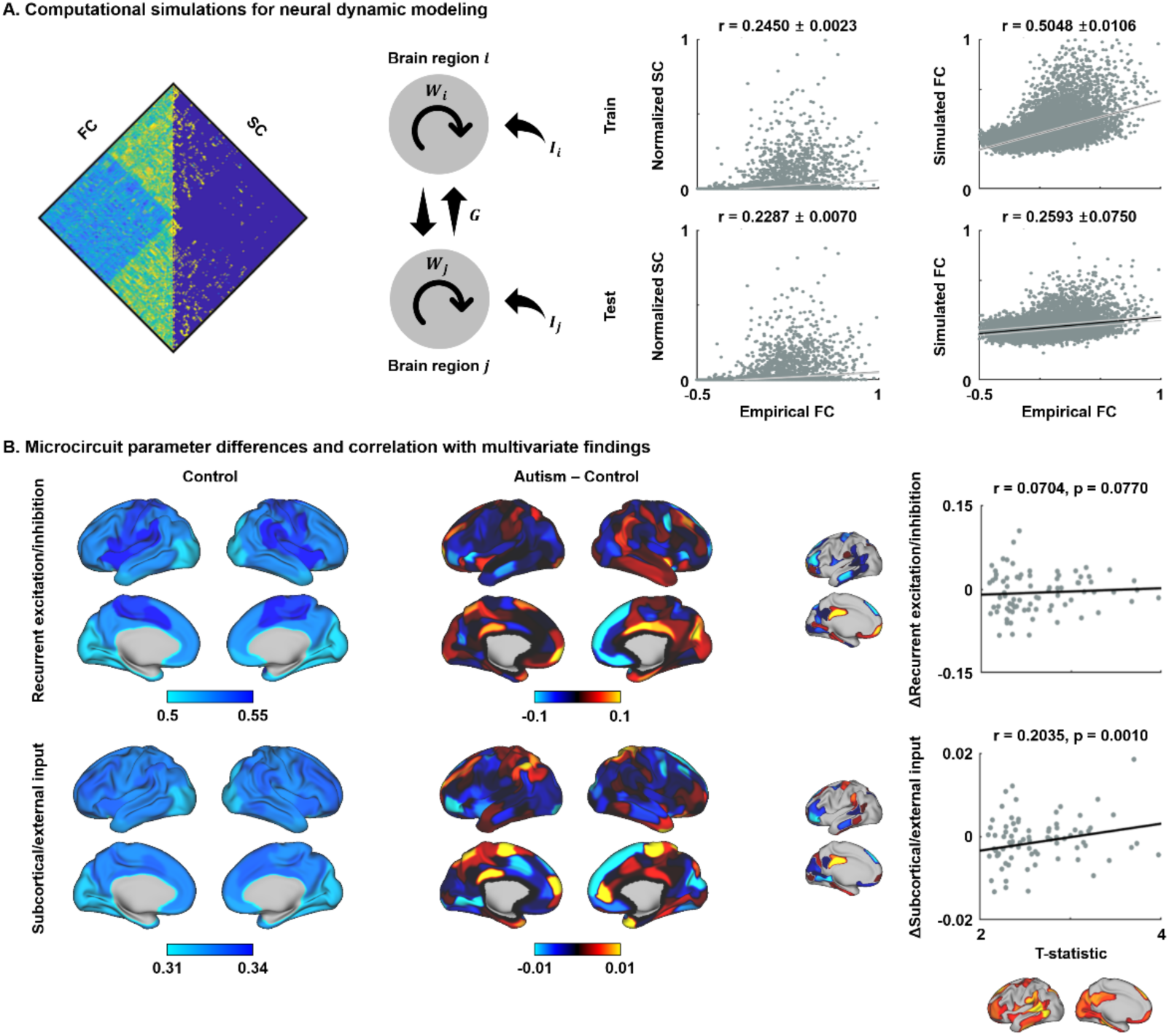
Microcircuit parameters and associations with macroscale findings. **(A)** A relaxed mean-field model was used to predict functional connectivity (FC) from structural connectivity (SC) and to estimate region specific microcircuit parameters *i*.*e*., recurrent excitation/inhibition *W*_*i*_ and subcortical/external input *I*_*i*_. A global coupling constant *G* is also estimated. Pearson’s correlations between FC and SC, and empirical and simulated FC are shown. Black lines indicate mean correlation and gray lines represent 95% confidence interval across cross-validation. **(B)** Microcircuit parameters of controls and differences of the parameters between individuals with autism and controls. Pearson’s correlations between t-statistics derived from the multivariate group comparison and the regional changes in microcircuit parameters are reported on the right side, constrained to regions showing significant between group differences in Fig 1.

Here, we harnessed a relaxed mean-field model ^18^ with five-fold cross-validation to first evaluate the capacity of the structural connectome to simulate intrinsic functional dynamics, and then estimated regional microcircuit parameters. The optimal model indeed predicted functional connectivity (Pearson’s correlation coefficient r ∼ 0.5 for training and r ∼ 0.26 for test data), outperforming baseline correlations between structural and functional connectivity (r ∼ 0.25 for training and r ∼ 0.23 for test data) (**Fig. 2A**). Although predictions were similar in autism and controls (**Fig. S5**), the model suggested distinct microcircuit configurations across groups (**Fig. 2B**). Compared to controls, individuals with autism displayed large differences in recurrent excitation/inhibition and subcortical input in heteromodal association and paralimbic areas followed by lower-level idiotypic and unimodal association areas. Correlating these regional changes in microcircuit patterns (see *Fig. 2B*) with multivariate macroscale manifold anomalies (see *Fig. 1C*), we observed a significant correlation between the overall degree of manifold distortion and increases in excitatory subcortical/external inputs (r = 0.2035 and p = 0.0010; determined using non-parametric spin tests that account for spatial autocorrelation ^39,58^) as well as marginal associations to increases in excitation/inhibition (r = 0.0704 and p = 0.0770) (**Fig. 2B**).

### Transcriptomic association analysis

We performed transcriptomic association analysis and developmental and disease enrichment analysis to explore potential neurobiological underpinnings of the above macroscale manifold findings (**Fig. 3A**). Specifically, we correlated the multivariate change pattern with post-mortem gene expression data from the Allen Institute for Brain Sciences (AIBS) ^59,60^. Significant gene lists after multiple comparisons correction (**Data S1**) were fed into a developmental gene expression analysis, which highlights developmental time windows across different brain regions in which these genes are strongly expressed (see *Methods*) ^51^. This analysis highlighted associations between the multivariate pattern of autism-related structural manifold distortions and genes expressed in early childhood and adolescence in thalamic as well as cortical areas (**Fig. 3B**). While these genes were also expressed in the cerebellum in later developmental stages, they were not significantly expressed in other subcortical regions such as the amygdala and striatum, nor in the hippocampus. In addition, we performed disease enrichment analysis to associate the significance of the gene expressions with the log fold-changes of autism, schizophrenia, and bipolar disorder (see *Methods*) ^61^. Notably, autism showed most marked associations (T = −34.89 and p < 0.001) followed by schizophrenia (T = −8.93 and p < 0.001) and bipolar disorder (T = 5.34 and p < 0.001) (**Fig. 3C**).

**Fig. 3.**
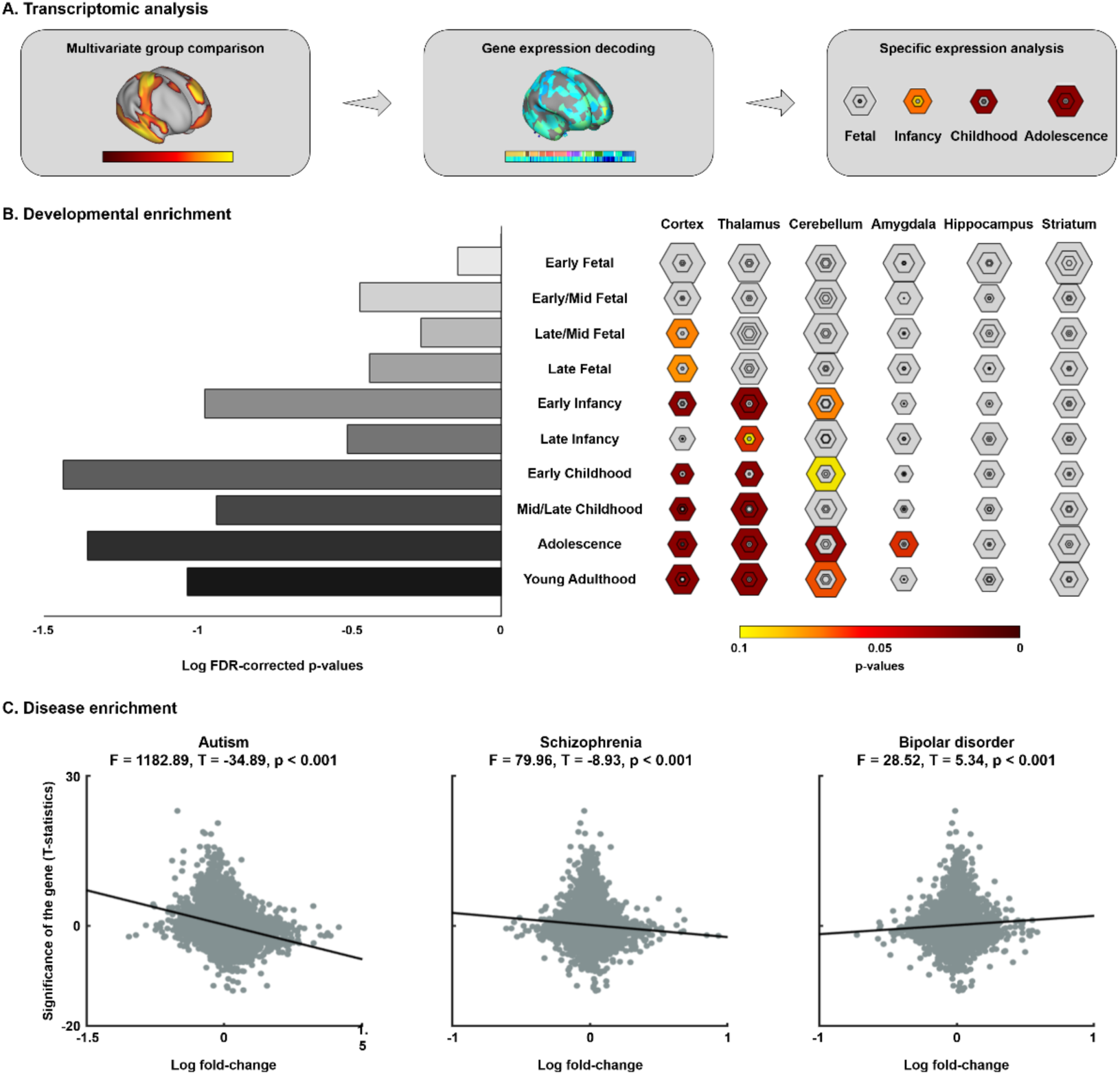
Transcriptomic analysis to identify gene expression patterns. **(A)** Multivariate findings of manifold distortions, gene expression decoding, and cell-type specific expression analysis. **(B)** Developmental enrichment, showing strong associations with cortex and thalamus during early childhood and adolescence. Hexagon rings represent the significance at different thresholds (from p < 0.05 in the outer ring to 0.0001 in the center). The bar plot on the left side represents the log transformed p-values that averaged across all brain structures that reported on the right side. **(C)** Disease enrichment analysis for associating gene expressions with disease effects of autism, schizophrenia, and bipolar disorder.

### Associations to symptom severity

We leveraged a supervised machine learning paradigm to predict symptom severity scores on the Autism Diagnostic Observation Schedule (ADOS – social cognition, communication, and repeated behavior/interest subscores and total score) ^62^ using structural connectome manifold information. Specifically, we employed elastic net regularization ^63^ with five-fold cross-validation (see *Methods*). The prediction procedure was performed 100 times with different set of training and test data to avoid subject selection bias. Structural connectome manifolds spanned by M1–M3 significantly predicted total ADOS score (mean ± SD r = 0.4182 ± 0.0806; mean ± SD mean absolute error (MAE) = 2.6899 ± 0.2215; permutation-test p < 0.03), as well as ADOS subscores of social cognition (mean ± SD r = 0.5295 ± 0.0620; mean ± SD MAE = 1.8195 ± 0.1604; permutation-test p < 0.003) and repeated behavior/interest subscores (mean ± SD r = 0.3269 ± 0.0856; mean ± SD MAE = 1.0007 ± 0.0847; permutation-test p < 0.09) (**Fig. 4**). On the other hand, manifold features did not predict ADOS subscores in the communication domain (mean ± SD r = 0.0409 ± 0.0939; mean ± SD MAE = 1.5876 ± 0.1755; and permutation-test p > 0.6). Selected features were primarily found in premotor/somatomotor areas, lateral and medial prefrontal cortices, as well as retrosplenial areas.

**Fig. 4.**
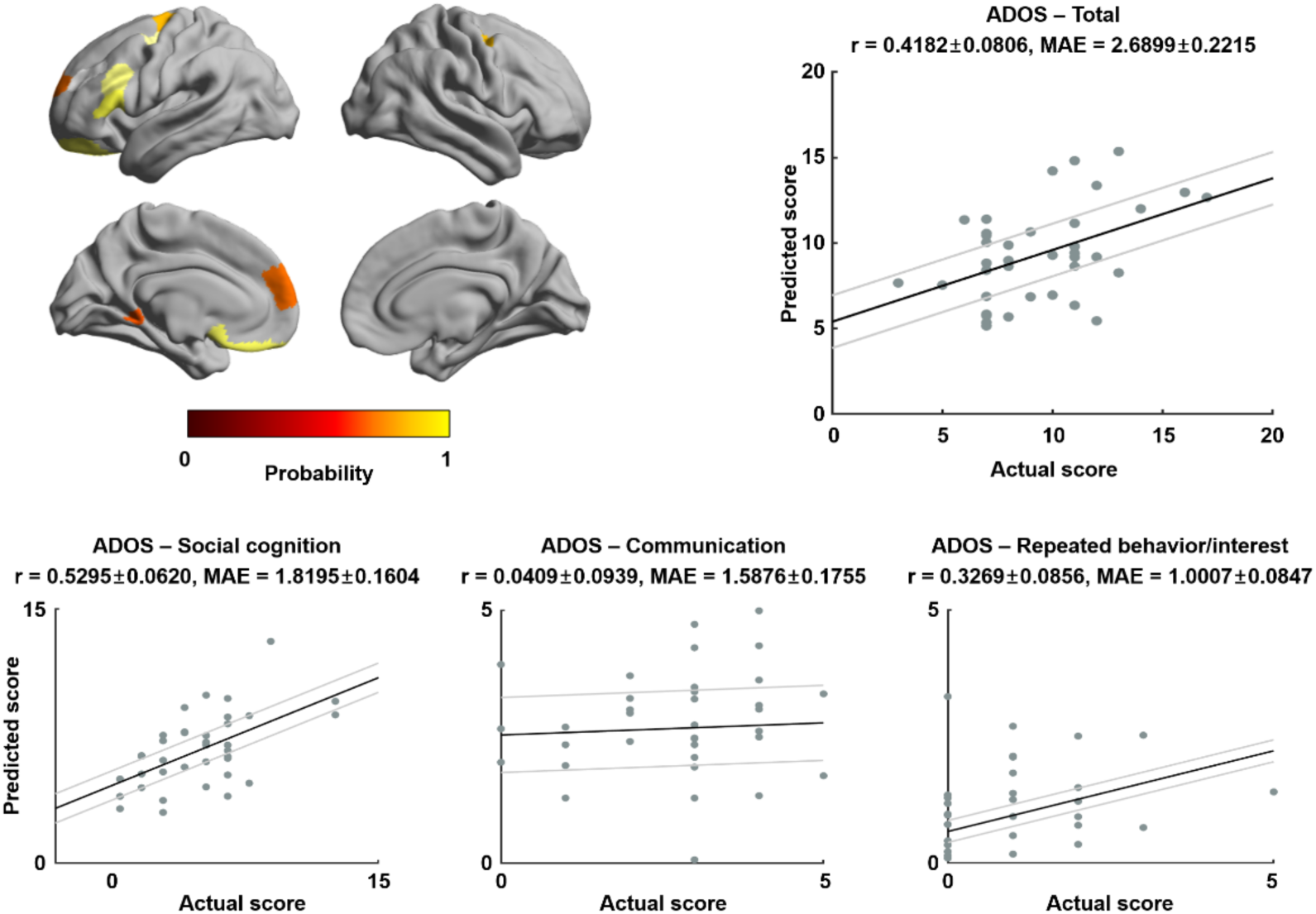
Associations between structural manifolds and autism symptoms. Probability of the selected brain regions across five-fold cross-validation for predicting ADOS scores is reported on the left top. Correlation between actual and predicted ADOS total and subscores are reported. Black line indicates mean correlation and gray lines represent 95% confidence interval for 100 iterations with different training/test dataset. *Abbreviation:* MAE, mean absolute error; ADOS, Autism Diagnostic Observation Schedule.

## Discussion

Understanding autism pathophysiology remains challenging, and a major complexity lies in difficulties to consolidate neuroimaging findings of connectome miswiring with molecular and neurophysiological data that focus on identifying cortical microcircuit changes. By combining novel manifold learning and computational models of brain dynamics, our study established how perturbations in macroscale structural connectome features in autism may relate to microcircuit dysfunction. We identified macroscale changes in cortical networks in autism, with peak differences in somatosensory as well as heteromodal association cortex, particularly the posterior core of the default-mode network. Findings were remained virtually unchanged when controlling for head motion and underlying morphological changes such as cortical thickness and curvature and were consistent across included study sites. Using biophysical parameters derived from a large-scale computational circuit model, we found that these whole-brain findings were correlated to alterations in subcortical drive into cortical microcircuits, together with marginal alterations in excitation/inhibition. An association to subcortical structures was also supported by complementary *post-mortem* transcriptomic association and developmental as well as disease enrichment analyses, highlighting that the affected regions harbor genes expressed in cortical and thalamic areas in early childhood and adolescence in autism. Our findings, therefore, offer a novel perspective on the relation between subcortico-cortical interactions at macroscale and microcircuit reorganization in autism.

The current work harnessed advanced manifold learning to compress high dimensional structural connectomes into a series of principal axes that describe spatial trends in connectivity changes across the cortical mantle in a data-driven manner. As such, our work brings a system level perspective to connectome reconfiguration in autism, which quantifies the integration and differentiation of brain subsystems in a continuous frame of reference. By offering a cortex-wide analysis of structural connectivity, our work extends prior diffusion MRI studies in autism that have shown atypical microstructure in fiber tracts underlying or interconnecting higher-order brain systems ^13,29,64–66^ and work focusing on fibers mediating connectivity between sensorimotor and subcortical systems ^7,67^. Our findings are also consistent with prior graph-theoretical studies of structural connectome data in autism that highlight alterations in global as well as local efficiency across both lower-level and higher-order cortical systems ^29,67–71^. In parallel, our findings provide new insights on potential structural substrates underlying a wide range of functional network anomalies reported in autism. ^6,72,73^. Functional findings are somewhat heterogenous across studies and analytical approaches; yet prior studies have converged on an overall pattern characterized by cortico-cortical functional connectivity reductions, often affecting in associative regions such as default-mode network, together with patches of connectivity increases, particularly between sensorimotor cortices and subcortical nodes such as the thalamus ^7,11,74,75^. By highlighting both association cortices such as the default mode network as well as idiotypic and somatosensory systems, our work provides a potential consolidation of these distributed effects in a space governed by structural wiring.

Histological studies have suggested several potential cellular substrates associated to connectome miswiring of autism including altered cortical lamination ^76–79^ and columnar layout ^80,81^, together with atypical neuronal migration that can result in cortical blurring ^77,82^ and changes in spine density of cortical projection neurons ^83,84^. Such cellular changes likely impact on the functional organization of cortical microcircuits in autism, also suggested by molecular studies in animals ^4,22–25,85,86^. These findings collectively support the imbalance in excitation and inhibition of cortical areas in autism ^4,9,16,22,87,88^, which have been related to anomalies in cortical neurotransmitter systems ^89–92^ and atypical subcortico-cortical interactions with subcortical structures such as thalamus modulating this balance ^7,11,24,74,75,93^. Here, we obtained support for perturbations in cortical microcircuit function from a network perspective, by leveraging a biophysically plausible computational model of brain function, which seeks to tune parameters to optimize the link between structural and connectomes ^18^. In recent work ^18^, these models were inverted to infer variations in the microcircuit level parameters at a cortical regional level. Studying a cohort of healthy adults from the human connectome project dataset, the study mapped cortex-wide gradients of recurrent excitation/inhibition and of excitatory subcortical input, with a topography mirroring prior work showing gradients in laminar differentiation and synaptic organization in non-human primates ^55,94^. In the neurotypical individuals studied here, we could show similar spatial trends in excitation/inhibition and subcortical input as in the prior work with overall increased subcortical input but lower excitation/inhibition in association cortices, while sensorimotor areas showed higher excitation/inhibition. This correspondence is an important consideration given that the retrospective data aggregated and shared via ABIDE is not at par in terms of image quality and data volume with the human connectome project data on which these models were originally presented ^18,95^. Importantly, comparing microcircuit maps between individuals with autism and controls supported overall a relatively diffuse pattern of local microcircuit parameter changes, not necessarily pointing to a unified direction of cortical microcircuit alterations. However, the connectome-wide manifold distortions were found to correlate with increases in excitatory subcortical drive into cortical microcircuits level, especially in lower-level cortical hierarchical areas, even after controlling for spatial autocorrelations using non-parametric spin tests, and to marginally relate to increases in excitation and inhibition.

Spatial decoding of macroscale distortion patterns with *post-mortem* gene expression maps provided potential etiological substrates of our findings. Recent studies in healthy brain organization ^96,97^, development ^37^, and disease ^98,99^ have shown that how such analyses can help understanding the relationship between macroscopic neuroimaging phenotypes and spatial variations at the molecular scale ^100^. In a prior study, similar approaches were used to identify genetic factors whose expression correlated to maps of cortical morphological variations in autism, and pointed to transcriptionally downregulated genes implicated in autism ^101^. In our study, similarly, developmental and disease enrichment analyses exhibited gene sets that were expressed in the cortex and thalamus during childhood and adolescence in autism by associating gene expression patterns with our macroscale findings, suggesting potential interactions between sensory-related subcortical areas and idiotypic/default-mode cortices. The thalamus relays afferent sensory inputs to the cortex and modulates efferent motor signals. Being a critical hub node in integrative cortico-cortical connectivity in both health and disease ^102,103^, the thalamus is furthermore recognized to regulate overall levels of cortical excitability ^7,11^. In other words, atypical excitatory input from thalamus would likely lead to altered perceptual input and may thus contribute to sensory abnormalities in autism ^11^. Indeed, it has been shown that abnormal functional connectivity between primary sensory cortices and subcortical regions affects the balance between sensory information processing and top-down feedback from higher-order cortices ^104^. Collectively, we can infer that the connectivity between thalamus and cortical areas extensively modulates brain-wide communication, which means abnormal thalamocortical connectivity likely affects multiple functional processes relevant to autism, including socio-cognitive impairments but also sensory anomalies ^7,11,24^.

Our multilevel analyses were carried out in a fully unconstrained manner, yet findings pointed to a co-existence of connectional and microcircuit perturbations in heteromodal association regions (particularly posterior default-mode nodes) and even more strongly to idiotypic/somatomotor cortices. In addition to the widely recognized impairments in communication and socio-cognitive functions, individuals with autism show obvious deficits in sensorimotor behaviors ^16,105^ and these are subsumed under the “repetitive behaviors and interests” syndrome cluster - a core criterion for autism diagnosis. More broadly, autism is increasingly thought to be associated to early sensory anomalies, also formulated in the so-called “sensory-first” hypothesis ^16^, where atypical formation and maturation of sensory processing circuits in early development results in a perturbed development of higher-order networks mediating more integrative and socio-communicative functions ^6,8^. Stratifying our cohort into children and adults, we also observed more marked sensorimotor network perturbations in the former, while anomalies in heteromodal association and paralimbic areas were only visible in adults with autism. Similar to recent work from our group that assessed functional hierarchy in autism ^6^, structural manifold features used in this study were particularly useful in predicting impairments in lower-level repetitive behavior symptoms as well as higher-order social and cognitive deficits, but not very sensitive in predicting atypical communication abilities. The inability of our approach to predict communication deficits remains to be investigated further, but it may potentially relate to other aspects of autism spectrum heterogeneity, including speech onset delay that affects only a subgroup of autism individuals and potentially associated compensatory mechanisms between sensory and language networks ^106–108^.

Our study provides a novel perspective consolidating brain organization at multiple scales to conceptualize autism pathophysiology. Harnessing advanced connectomics, machine learning, and computational modeling, we could show macroscale structural connectome perturbations in somatosensory/idiotypic and default-mode/heteromodal association areas in autism, which are associated with behavioral symptoms at an individual subject level. These macroscale distortions were also found to relate to cortical microcircuit function in individuals with autism in an *in silico* model of brain function, in our cohort mostly visible as an increase in excitatory subcortical drive. Arguably, the modest sample size of this study warrants further replication of our findings in large, and ideally transdiagnostic cohorts, to also evaluate specificity of our findings and to further explore heterogeneity within the autism spectrum itself ^106,107,109,110^. Yet, these findings overall provide consistent support that atypical subcortico-cortical interactions, likely between thalamic and sensorimotor areas, contribute to large-scale network anomalies in autism and may suggest that connectivity anomalies of sensorimotor networks that mature early may cascade into an overall disorganization of cortico-cortical systems in autism.

## Methods

### Participants

We studied imaging and phenotypic data of 47 individuals with autism and 37 typically developing controls from the Autism Brain Imaging Data Exchange initiative (ABIDE-II; http://fcon_1000.projects.nitrc.org/indi/abide) ^52,53^. Participants were taken from two independent sites: (1) New York University Langone Medical Center (NYU) and (2) Trinity College Dublin (TCD), which were the only sites that included children and adults with autism and neurotypical controls, with ≥10 individuals per group, and who had full MRI data (*i*.*e*. structural, functional, and diffusion) available. These 84 participants were selected from a total of 120 participants through the following inclusion criteria: (i) complete multimodal imaging data *i*.*e*., T1-weighted, resting-state functional MRI (rs-fMRI), and dMRI, (ii) acceptable cortical surface extraction, (iii) low head motion in the rs-fMRI time series *i*.*e*., less than 0.3 mm framewise displacement. Individuals with autism were diagnosed by an in-person interview with clinical experts and gold standard diagnostics of Autism Diagnostic Observation Schedule (ADOS) ^62^ and/or Autism Diagnostic Interview-Revised (ADI-R) ^111^. Neurotypical controls did not have any history of mental disorders. For all groups, participants who had genetic disorders associated with autism (*i*.*e*., Fragile X), psychological disorders comorbid with autism, contraindications to MRI scanning, and pregnant were excluded. The ABIDE data collections were performed in accordance with local Institutional Review Board guidelines, and data were fully anonymized. Detailed demographic information of the participants is reported in **Table S1**.

### MRI acquisition

At the NYU site, multimodal imaging data were acquired using 3T Siemens Allegra. T1-weithed data were obtained using a 3D magnetization prepared rapid acquisition gradient echo (MPRAGE) sequence (repetition time (TR) = 2,530 ms; echo time (TE) = 3.25 ms; inversion time (TI) = 1,100 ms; flip angle = 7°; matrix = 256 × 192; and voxel size = 1.3 × 1.0 × 1.3 mm^3^). The rs-fMRI data were acquired using a 2D echo planar imaging (EPI) sequence (TR = 2,000 ms; TE = 15 ms; flip angle = 90°; matrix = 80 × 80; number of volumes = 180; and voxel size = 3.0 × 3.0 × 4.0 mm^3^). Finally, dMRI data were obtained using a 2D spin echo EPI (SE-EPI) sequence (TR = 5,200 ms; TE = 78 ms; matrix = 64 × 64; voxel size = 3 mm^3^ isotropic; 64 directions; b-value = 1,000 s/mm^2^; and 1 b0 image).

At the TCD site, imaging data were acquired using 3T Philips Achieva. T1-weighted MRI were acquired using a 3D MPRAGE (TR = 8,400 ms; TE = 3.90 ms; TI = 1,150 ms; flip angle = 8°; matrix = 256 × 256; voxel size = 0.9 mm^3^ isotropic). The rs-fMRI data were aquired using a 2D EPI (TR = 2,000 ms; TE = 27 ms; flip angle = 90°; matrix = 80 × 80; number of volumes = 210; and voxel size = 3.0 × 3.0 × 3.2 mm^3^). Finally, dMRI data were acquired using a 2D SE-EPI (TR = 20,244 ms; TE = 79 ms; matrix = 124 × 124; voxel size = 1.94 × 1.94 × 2 mm^3^; 61 directions; b-value = 1,500 s/mm^2^; and 1 number b0 image).

### Data preprocessing

T1-weighted data were processed using FreeSurfer ^112–117^, which includes gradient nonuniformity correction, skull stripping, intensity normalization, and tissue segmentation. White and pial surfaces were generated through triangular surface tessellation, topology correction, inflation, and spherical registration to fsaverage. Rs-fMRI data were processed via C-PAC (https://fcp-indi.github.io) ^118^, including slice timing and head motion correction, skull stripping, and intensity normalization. Nuisance variables of head motion, average white matter and cerebrospinal fluid signal, and linear/quadratic trends were removed using CompCor ^119^. Band-pass filtering between 0.01 and 0.1 Hz was applied, and rs-fMRI data were co-registered to T1-weighted data in MNI standard space with linear and non-linear transformations. The rs-fMRI data were mapped to subject-specific mid-thickness surfaces and resampled to the Conte69 template. Finally, surface-based spatial smoothing with a full width at half maximum of 5 mm was applied. The dMRI data was processed using MRtrix ^30,31^ including correction for susceptibility distortions, head motion, and eddy currents. Quality control involved visual inspection of T1-weighted data, and cases with faulty cortical segmentation were excluded. Data with a framewise displacement of rs-fMRI data >0.3 mm were also excluded ^120,121^.

### Structural connectome manifold identification

Structural connectomes were generated from preprocessed dMRI data using MRtrix ^30,31^. Anatomical constrained tractography was performed using different tissue types derived from the T1-weighted image, including cortical and subcortical grey matter, white matter, and cerebrospinal fluid ^33^. The multi-shell and multi-tissue response functions were estimated ^35^ and constrained spherical-deconvolution and intensity normalization were performed ^34^. The tractogram was generated with 40 million streamlines, with a maximum tract length of 250 and a fractional anisotropy cutoff of 0.06. Subsequently, spherical-deconvolution informed filtering of tractograms (SIFT2) was applied to reconstruct whole-brain streamlines weighted by the cross-section multipliers ^32^. The structural connectome was built by mapping the reconstructed cross-section streamlines onto the Schaefer atlas with 200 parcels ^32,122^ then log-transformed ^123^.

Cortex-wide structural connectome manifolds were identified using BrainSpace (https://github.com/MICA-MNI/BrainSpace) ^39^. First, a template manifold was estimated using a group representative structural connectome, defined using a distance-dependent thresholding that preserves long-range connections ^54^. The group representative structural connectome was constructed using both autism and control data. A cosine similarity matrix, capturing similarity of connections among different brain regions, was constructed without thresholding the structural connectome and manifolds were estimated via diffusion map embedding (**Fig. 1A–B**). Diffusion map embedding is robust to noise and computationally efficient compared to other non-linear manifold learning techniques ^124,125^. It is controlled by two parameters α and t, where α controls the influence of the density of sampling points on the manifold (α = 0, maximal influence; α = 1, no influence) and t controls the scale of eigenvalues of the diffusion operator. We set α = 0.5 and t = 0 to retain the global relations between data points in the embedded space, following prior applications ^6,38,39,48^. In this new manifold, interconnected brain regions are closely located and regions with weak inter-connectivity located farther apart. After generating the template manifold, individual-level manifolds were then estimated and aligned to the template manifold via Procrustes alignment.

### Between-group differences in structural manifolds

Multivariate analyses compared individuals with autism and controls in the manifold spanned by the first three structural eigenvectors, which explained more than 50% in structural connectome variance and corresponded to the elbow in the scree plot. Models controlled for age, sex, and site in addition to including the group factor. We corrected for multiple comparisons using the FDR procedure ^126^. Summary statistics were calculated based on an atlas of laminar differentiation and cortical hierarchy (**Fig. 1C**) ^55^ and a widely used community parcellation (**Fig. S1A**) ^56^. To simplify the multivariate manifold representations into a single scalar, we quantified manifold distance as the Euclidean distance between the center of template manifold and all data points (*i*.*e*., brain regions) in the manifold space for each individual after alignment (**Fig. S1B**) ^57^. Group averaged manifold distance was compared between individuals with autism and controls to assess the manifold-affected brain regions.

### Microscale neural dynamic modeling

A large-scale biophysical dynamic circuit modeling was conducted to predict functional connectivity from structural connectome information and to estimate regional microcircuit parameters. Specifically, we harnessed a relaxed mean-field model that captures the link between cortical functional dynamics and structural connectivity derived from dMRI, and its modulation through region-specific microcircuit parameters ^18^. In comparison to other models that also include synapse-level parameters, this model has a more synoptic scale, allowing for structure-function simulations with modest parametric complexity. For details on the model and its mathematical underpinnings, we refer to the original publication on the relaxed mean-field model ^18^ and earlier work on the use of (non-relaxed) mean-field models ^40^. In brief, these models approximate the dynamics of spiking and interconnected neural networks through a simplified one-dimensional equation. Mean-field models assume that neural dynamics of a given region are governed by (i) recurrent intra-regional input *i*.*e*., recurrent excitation/inhibition; (ii) inter-regional input, mediated by dMRI-based structural connections from other nodes, (iii) extrinsic input, mainly from subcortical regions, and (iv) neuronal noise ^18^. While the original (non-relaxed) mean-field models ^40^ assume these parameters to be constant across brain regions, the relaxed mean-field variant allows recurrent excitation/inhibition and subcortical/external input to vary. In the model, global brain dynamics of the network of interconnected local networks is described by the following three coupled nonlinear stochastic differential equations ^18^:

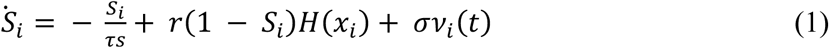

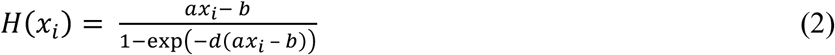

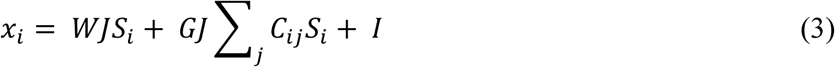

For a given region *i, S*_*i*_ in formula (1) represents the average synaptic gating variable, *H(x*_*i*_) in formula (2) is the population firing rate, and *x*_*i*_ in formula (3) is the total input current. The input current *x*_*i*_ is determined by the recurrent connection strength *W*_*i*_ (*i*.*e*., *excitation/inhibition*) and the excitatory input *I*_*i*_, such as from subcortical relays (*i*.*e*., *subcortical/external input*), and inter-regional signal flow. The latter is governed by *C*_*ij*_, which represents the structural connectivity between regions *i* and *j*, and the global coupling *G*. In equation (1), the *v*_*i*_ term refers to uncorrelated Gaussian noise, modulated by an overall noise amplitude *σ*. Following prior work ^18^, we set parameters as *J* = 0.2609 nA, *a* = 270 *n/C, b* = 108 Hz, *d* = 0.154 s, *r* = 0.641, and *τs* = 0.1s.

We fed the group representative structural connectivity matrix and group averaged functional connectivity matrix into the relaxed mean-field model optimization, which provided recurrent connection strengths *W* and excitatory subcortical inputs *I* for every cortical region, as well as a global coupling constant *G* and global noise amplitude *σ*. Global and region-specific parameters were determined by maximizing the similarity between simulated and empirical functional connectivity, based on a previously developed algorithm for inverting neural mass models that leverages the well-established expectation-maximization algorithm ^127,128^. Pearson’s correlations between empirical functional connectivity and structural connectivity, and that with the simulated functional connectivity of control data were calculated to assess the quality of the microcircuit parameter estimation (**Fig. 2A**). These procedures were performed with a five-fold cross-validation framework with random separation of training and test data, and final microcircuit parameters were determined by averaging across cross-validations.

The estimated recurrent excitation/inhibition and subcortical/external input parameters were compared between individuals with autism and controls. Pearson’s correlations were calculated between the differences in these model-derived parameters between groups and t-statistics of multivariate analysis (**Fig. 2B**) to evaluate the association between macroscale structural connectome reorganization and imbalances in microcircuit properties. Significances of spatial correlations were assessed via 1,000 spin test permutations with randomly rotated microcircuit parameters ^58^.

### Transcriptomic association analysis

To provide additional neurobiological context of our findings, we assessed spatial correlations between the between-group differences in the structural manifold and gene expression patterns (**Fig. 3A**). Initially, we correlated the t-statistics map derived from the multivariate group comparison and the post-mortem gene expression maps provided by Allen Institute for Brain Sciences (AIBS) using the Neurovault gene decoding tool ^59,60^. Neurovault implements mixed-effect analysis to estimate associations between the input t-statistic map and the genes of AIBS donor brains yielding the gene symbols associated with the input t-statistic map. Gene symbols that passed for a significance level of FDR-corrected p < 0.05 were considered for the subsequent analysis. In a second stage, gene lists that were significant were fed into enrichment analysis (**Fig. 3B**), which involved comparison against developmental expression profiles from the BrainSpan dataset (http://www.brainspan.org) using the cell-type specific expression analysis (CSEA) developmental expression tool (http://genetics.wustl.edu/jdlab/csea-tool-2) ^51^. As the AIBS repository is composed of adult post-mortem datasets, it should be noted that the associated gene symbols represent indirect associations with the input t-statistic map derived from the developmental data. To explore whether the Neurovault derived genetic signature was associated with autism pathophysiology, we additionally performed disease enrichment analysis using previously published transcriptome findings for autism, schizophrenia, and bipolar disorder (**Fig. 3C**) ^61^. A robust linear regression model was constructed for linking the significance of the gene expressions (*i*.*e*., t-statistic) derived from Neurovault with log fold-change of autism, schizophrenia, and bipolar disorder, which share similar genetic variants ^129^. The fold-change represents the level to which a gene is over or under expressed in a particular condition ^61^. Guanine-cytosine (GC) content was controlled to avoid possible effects that related to genome size in microarray data ^130,131^.

### Symptom severity prediction

We adopted a supervised machine learning framework with cross-validation to predict autism symptoms as measured by ADOS scores ^62^. We aimed at predicting total ADOS scores, as well as subscores for social cognition, communication, and repeated behavior/interest (**Fig. 4**). We utilized five-fold cross-validation separating training and test data. Feature selection procedure was conducted using the training data (4/5 segments) and it was repeated 5 times with different segments of training data. Among a total of 600 (200 regions × 3 gradients) features, a set of features that could predict each ADOS score was identified using elastic net regularization (regularization parameter = 0.5), which shows good performance in feature selection at a given sparsity level compared to L1 (least absolute shrinkage and selection operator)- and L2 (ridge)-norm regularization methods ^63^. The features that were most frequently identified across the cross-validation iterations (>50%) were selected to predict each ADOS score. Linear regression model for predicting ADOS scores was constructed using the selected features controlled for age, sex, and site as independent variables within the training data (4/5 segments) and it was applied to the test data (1/5 segment) to predict their ADOS scores. Prediction procedure was repeated 100 times with different set of training and test data to avoid bias for separating subjects. The prediction accuracy was assessed by calculating Pearson’s correlation between the actual and predicted ADOS scores as well as their mean absolute error (MAE). 95% confidence interval of accuracy measures were also reported. Permutation-based correlations across 1,000 spin tests were conducted by randomly shuffling ADOS scores to check whether the prediction performance exceeded chance levels ^58^.

### Sensitivity and specificity analyses

#### a) Site effects

The multivariate group comparison using structural connectome manifolds was performed for each site (NYU and TCD separately) to see the consistency of results across different sites (**Fig. S1C**).

#### b) Head motion effects

To rule out whether the macroscale perturbations in autism related to head motions, we first calculated mean FD from dMRI for all participants (**Fig. S2A**). Two-sample t-test then assessed between-group differences in head motion. In addition, we repeated the multivariate manifold comparisons between groups while controlling for mean FD (**Fig. S2B**).

#### c) Age effects

To assess the age-related effects on structural connectome manifolds, we performed multivariate group comparison in manifolds, controlled for sex and site, within children (age < 18) and adults (age ≥ 18) cohorts separately (**Fig. S3A**).

#### d) Associations to cortical morphology

Several studies have previously reported atypical cortical morphology in individuals with autism relative to controls ^12,19^. To assess whether these morphological variations contribute to our connectome results, we calculated Pearson correlations between multivariate findings and cortical morphology measures (*i*.*e*., cortical thickness and cortical curvature) between groups (**Fig. S4A**). We also repeated the multivariate manifold comparisons while controlling for cortical thickness and curvature, to evaluate whether the connectome-wide effects can be observed above and beyond potential variations in cortical morphology (**Fig. S4B**).

## Acknowledgements

Dr Bo-yong Park was funded by Molson Neuro-Engineering fellowship by Montreal Neurological Institute and Hospital (MNI). Dr Casey Paquola was funded through a postdoctoral fellowship of the Fonds de la Recherche due Quebec – Santé (FRQ-S). Dr Oualid Benkarim was funded by a Healthy Brains for Healthy Lives (HBHL) postdoctoral fellowship. Dr Richard A. I. Bethlehem was funded by a British Academy Post-Doctoral Fellowship and the Autism Research Trust. Dr B.T. Thomas Yeo was supported by the Singapore National Research Foundation (NRF) Fellowship (Class of 2017). Any opinions, findings and conclusions or recommendations expressed in this material are those of the authors and do not reflect the views of National Research Foundation, Singapore. Dr Jonathan Smallwood was supported by the European Research Council (WANDERINGMINDS-ERC646927). Dr Boris C. Bernhardt acknowledges research support from the National Science and Engineering Research Council of Canada (NSERC Discovery-1304413), the Canadian Institutes of Health Research (CIHR FDN-154298), SickKids Foundation (NI17-039), Azrieli Center for Autism Research (ACAR-TACC), and the Tier-2 Canada Research Chairs program. Drs Bo-yong Park, Casey Paquola, Richard A. I. Bethlehem, and Boris C. Bernhardt are jointly funded through an MNI-Cambridge collaborative award.

## Author contributions

B.P. and B.C.B. designed the experiments, analyzed the data, and wrote the manuscript. S.H., C.P. and O.B. aided with the experiments. S.V., R.A.I.B, A.D.M., M.P.M., A.G., B.T.T.Y., L.M., and J.S. reviewed the manuscript. B.P. and B.C.B. are the corresponding authors of this work and have responsibility for the integrity of the data analysis.

