## Supplementary Materials for "Connectome and microcircuit models implicate atypical subcortico-cortical interactions in autism pathophysiology"

*[a]McConnell Brain Imaging Centre, Montreal Neurological Institute and Hospital, McGill University, Montreal, Quebec, Canada; [b]Center for the Developing Brain, Child Mind Institute, New York City, New York, United States of America; [c]Forschungszentrum Julich, Germany; [d]Max Planck Institute for Cognitive and Brain Sciences, Leipzig, Germany; [e]Autism Research Centre, Department of Psychiatry, University of Cambridge, Cambridge, United Kingdom; [f]Brain Mapping Unit, Department of Psychiatry, University of Cambridge, Cambridge, United Kingdom; [g]Functional Neuroimaging Laboratory, Istituto Italiano di Tecnologia, Centre for Neuroscience and Cognitive Systems @ UNITN, Rovereto, Italy; [h]Department of Electrical and Computer Engineering, Centre for Sleep and Cognition, Clinical Imaging Research Centre and N.1 Institute for Health, National University of Singapore, Singapore; [i]Martinos Center for Biomedical Imaging, Massachusetts General Hospital, Charlestown, Massachusetts, United States of America; [j]NUS Graduate School for Integrative Sciences and Engineering, National University of Singapore, Singapore; [k]Department of Psychology, York Neuroimaging Centre, University of York, York, United Kingdom*

***Corresponding Authors:**

Boris C. Bernhardt, PhD
Multimodal Imaging and Connectome Analysis Lab
McConnell Brain Imaging Centre
Montreal Neurological Institute and Hospital
McGill University
Montreal, Quebec, Canada


Bo-yong Park, PhD
Multimodal Imaging and Connectome Analysis Lab
McConnell Brain Imaging Centre
Montreal Neurological Institute and Hospital
McGill University
Montreal, Quebec, Canada


**Table S1 | Demographic information of the study participants.** Means and SDs are reported.

| Information | | NYU | | TCD | | P-value |
|---|---|---|---|---|---|---|
| Number (Autism/Control) | | 29/18 | | 18/19 | | 0.2316* |
| Age | Autism | 9.61 (6.16) | p = 0.8074 | 14.46 (3.30) | p = 0.2088 | 0.0037 |
| | Control | 10.01 (3.95) | | 15.83 (3.21) | | <0.001 |
| Sex (male:female) | Autism | 24:5 | p = 0.2432* | 18:0 | p = 1* | 0.0624* |
| | Control | 17:1 | | 19:0 | | 0.2976* |
| ADOS – Total | | 10.00 (3.36) | | 8.72 (2.44) | | 0.1924 |
| ADOS – Social cognition | | 7.50 (2.09) | | 5.78 (2.37) | | 0.0226 |
| ADOS – Communication | | 2.5 (1.70) | | 2.94 (0.87) | | 0.3260 |
| ADOS – Repeated behavior/interest | | 1.40 (1.27) | | 0.22 (0.55) | | <0.001 |

*Chi-squared

*Abbreviations:* SD, standard deviation; NYU, New York University Langone Medical Center; TCD, Trinity College Dublin; ADOS, Autism Diagnostic Observation Schedule.

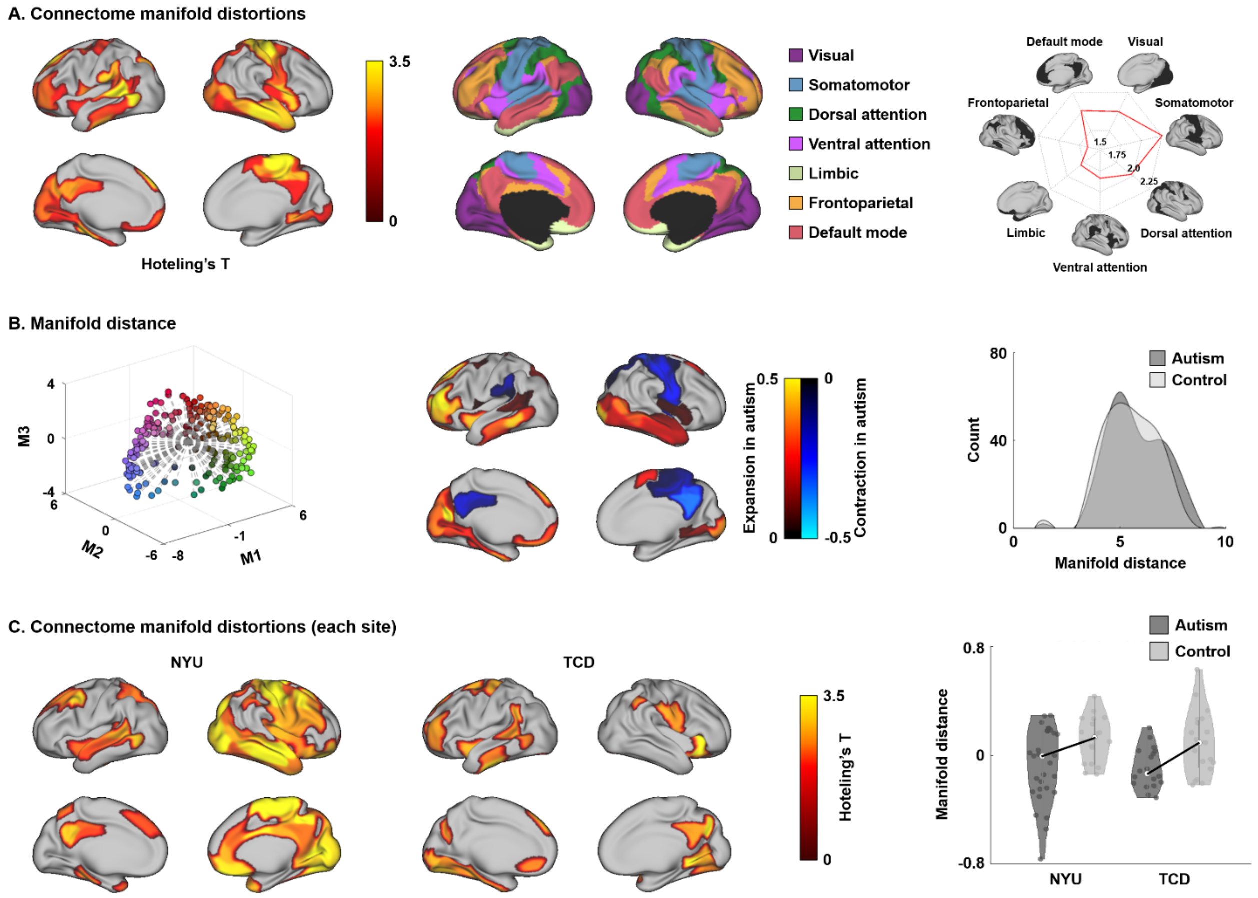


**Fig. S1 | Connectome manifold distortions. (A)** The t-statistics of the identified regions that showed significant differences in manifolds between autism and controls. Findings were corrected for multiple comparisons at FDR < 0.05. Functional network-wise [54] summary of the t-statistic values are shown in radar plots. **(B)** Manifold distance measured as the Euclidean distance between the template center and each data point and its difference between the groups. The histogram of the manifold distance for each group is reported. **(C)** The t-statistics derived from the multivariate group comparison for each site. Manifold distance perturbations in autism and controls of the identified regions for each site are reported on the right side. *Abbreviations:* FDR, false discovery rate; NYU, New York University Langone Medical Center; TCD, Trinity College Dublin.

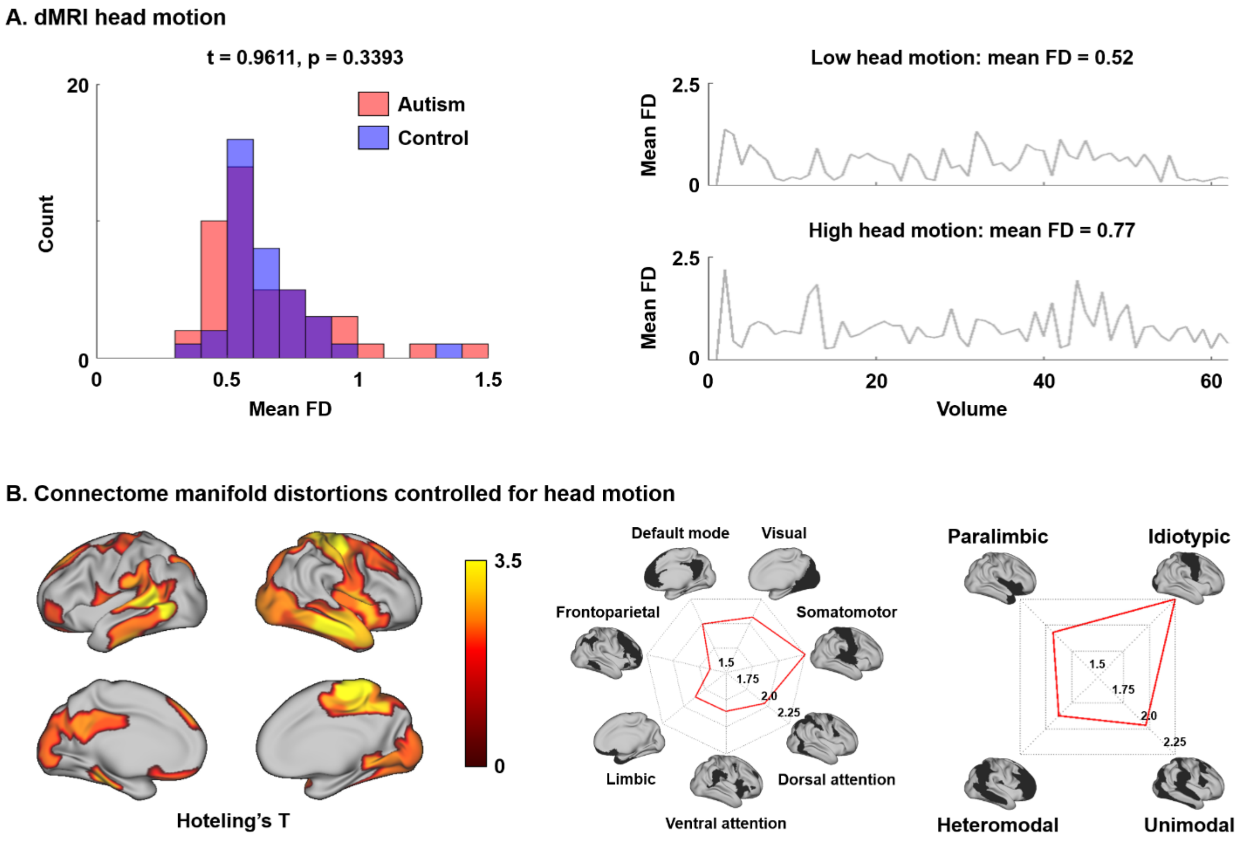


**Fig. S2 | Head motion effects. (A)** Mean framewise displacement (FD) of each group and that of two representative participants with low and high head motion. **(B)** The t-statistics derived from the multivariate group comparison between autism and controls using the three structural manifolds controlled for head motion.

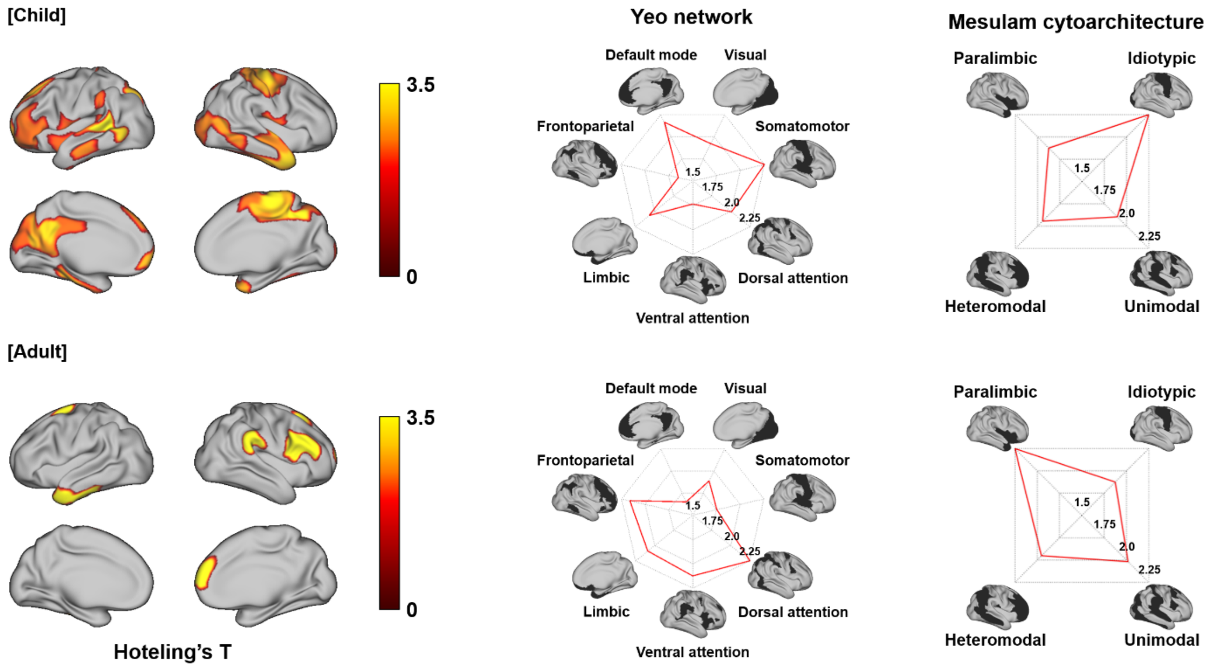


**Fig. S3 | Age effects.** The t-statistics derived from the multivariate group comparison between autism and controls using the three structural manifolds within children and adult cohorts, separately.

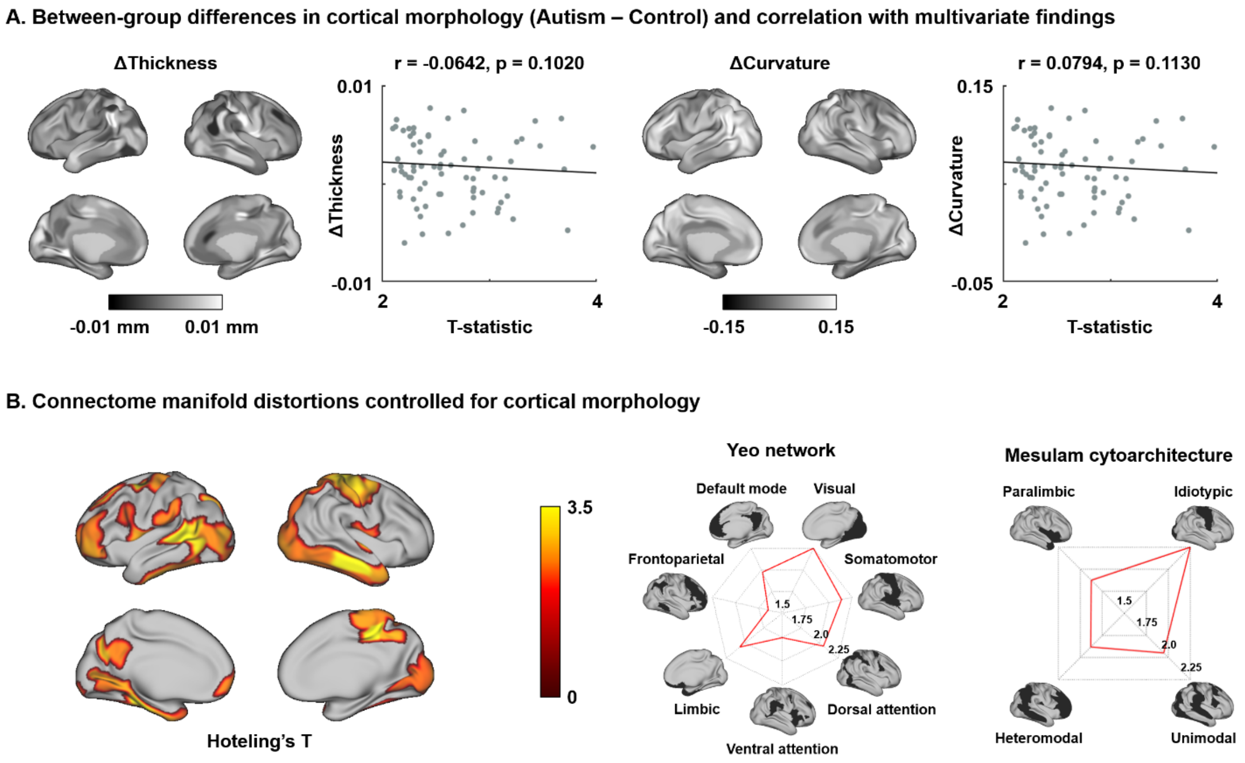


**Fig. S4 | Morphological associations. (A)** Correlation between differences in cortical morphology (*i.e.* thickness and folding) between autism and controls and multivariate change pattern. **(B)** The t-statistics derived from the multivariate group comparison between individuals with autism and controls using the three structural manifolds controlled for cortical morphology.

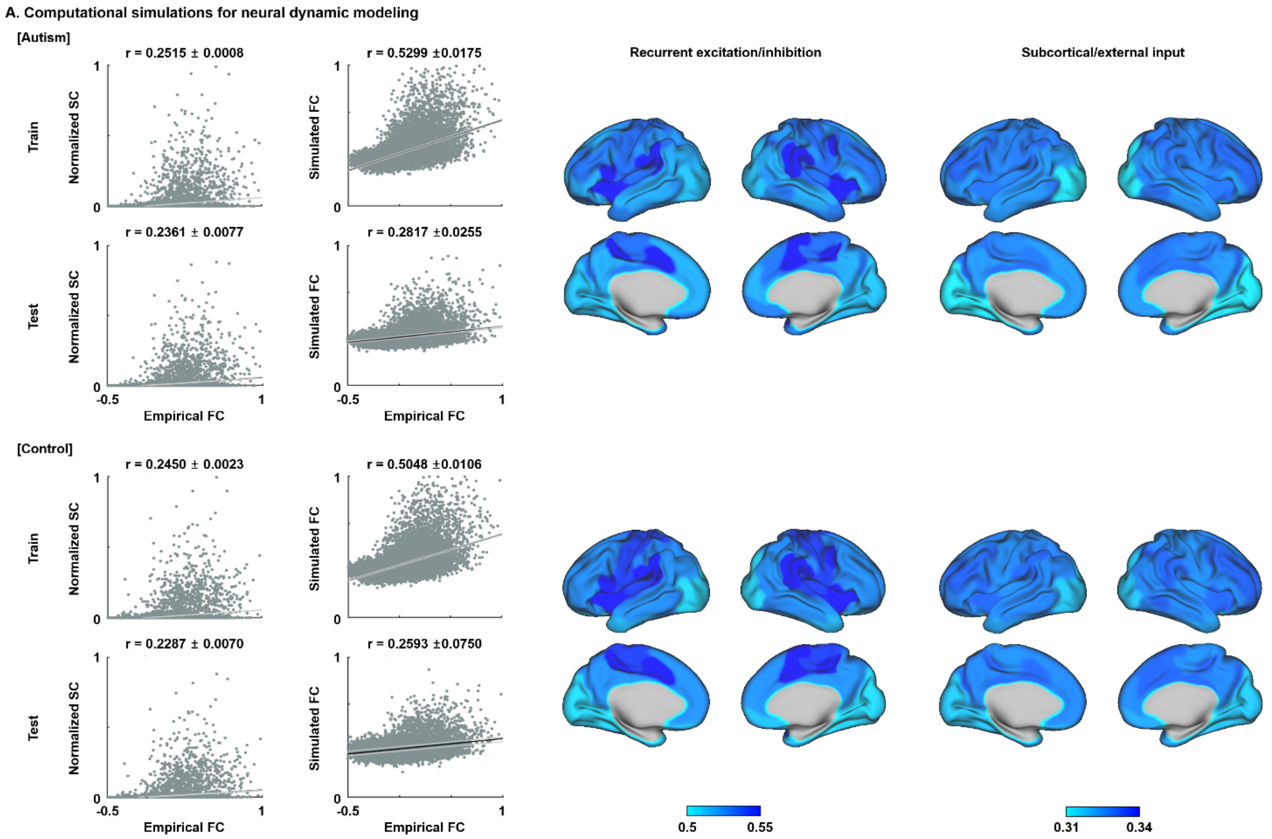


**Fig. S5 | Biophysical simulations.** Pearson's correlations between functional connectivity (FC) and structural connectivity (SC), and empirical and simulated FC, as well as microcircuit parameters for individuals with autism and controls. Black lines indicate mean correlation and gray lines represent 95% confidence interval across cross-validation.

**Data S1 | Significant gene lists correlated with multivariate change pattern.** Gene symbol with name and t-statistic as well as false discover rate corrected p-value are reported in the Supplementary Data file (Supplementary_Data1.xlsx).

| Symbol | Name | t | p-FDR |
|---|---|---|---|
| CBLN4 | cerebellin 4 precursor | 54.15 | 0.001 |
| THEG | theg spermatid protein | 22.96 | 0.027 |
| COL5A1 | collagen, type V, alpha 1 | 20.52 | 0.027 |
| PLXDC1 | plexin domain containing 1 | 18.76 | 0.027 |
| PRSS16 | protease, serine, 16 (thymus) | 18.45 | 0.027 |
| ADPRHL1 | ADP-ribosylhydrolase like 1 | 18.33 | 0.027 |
| CHRNA2 | cholinergic receptor, nicotinic, alpha 2 (neuronal) | 15.82 | 0.030 |
| TRIM54 | tripartite motif containing 54 | 15.8 | 0.030 |
| RORB | RAR-related orphan receptor B | 15.71 | 0.030 |
| CCNO | cyclin O | 15.75 | 0.030 |
| MYO15A | myosin XVA | 15.17 | 0.030 |
| NOS2 | nitric oxide synthase 2, inducible | 14.98 | 0.030 |
| STX19 | syntaxin 19 | 14.85 | 0.030 |
| CUX2 | cut-like homeobox 2 | 14.63 | 0.030 |
| EIF4E1B | eukaryotic translation initiation factor 4E family member 1B | 14.36 | 0.030 |
| SORCS1 | sortilin-related VPS10 domain containing receptor 1 | 14.21 | 0.030 |
| ABLIM2 | actin binding LIM protein family, member 2 | 13.94 | 0.030 |
| CAMK2G | calcium/calmodulin-dependent protein kinase II gamma | 13.87 | 0.030 |
| RGS6 | regulator of G-protein signaling 6 | 13.89 | 0.030 |
| REEP5 | receptor accessory protein 5 | 13.71 | 0.030 |
| NLRP3 | NLR family, pyrin domain containing 3 | 13.68 | 0.030 |
| LTK | leukocyte receptor tyrosine kinase | 13.42 | 0.030 |
| KCNJ3 | potassium inwardly-rectifying channel, subfamily J, member | 13.35 | 0.030 |
| IFT122 | intraflagellar transport 122 homolog (Chlamydomonas) | 13.31 | 0.030 |
| IFIT1 | interferon-induced protein with tetratricopeptide repeats 1 | 13.2 | 0.030 |
| SNAP25 | synaptosomal-associated protein, 25kDa | 13.12 | 0.030 |
| FBLN7 | fibulin 7 | 13.03 | 0.030 |
| NKRF | NFKB repressing factor | 12.63 | 0.030 |
| COX7A1 | cytochrome c oxidase subunit VIIa polypeptide 1 (muscle) | 12.6 | 0.030 |
| AMIGO1 | adhesion molecule with Ig-like domain 1 | 12.61 | 0.030 |
| DBH | dopamine beta-hydroxylase (dopamine beta- | 12.54 | 0.030 |
| C16orf58 | chromosome 16 open reading frame 58 | 12.52 | 0.030 |
| VSNL1 | visinin-like 1 | 12.44 | 0.030 |
| CTNNAL1 | catenin (cadherin-associated protein), alpha-like 1 | 12.47 | 0.030 |
| CYP26B1 | cytochrome P450, family 26, subfamily B, polypeptide 1 | 12.18 | 0.031 |
| MGAT5B | mannosyl (alpha-1,6-)-glycoprotein beta-1,6-N-acetyl-glucosaminyltransferase, isozyme B | 12.01 | 0.031 |
| UNC5D | unc-5 homolog D (C. elegans) | 11.97 | 0.031 |
| ZMAT4 | zinc finger, matrin-type 4 | 11.76 | 0.032 |
| LAMA3 | laminin, alpha 3 | 11.64 | 0.032 |
| ANKRD24 | ankyrin repeat domain 24 | 11.58 | 0.032 |
| SMYD2 | SET and MYND domain containing 2 | 11.57 | 0.032 |
| PKNOX2 | PBX/knotted 1 homeobox 2 | 11.54 | 0.032 |
| GNB1L | guanine nucleotide binding protein (G protein), beta | 11.2 | 0.036 |
| TMEM38A | transmembrane protein 38A | 11.2 | 0.036 |
| STAMBPL1 | STAM binding protein-like 1 | 11.11 | 0.036 |
| KCNS1 | potassium voltage-gated channel, delayed-rectifier, | 10.95 | 0.037 |
| VIPR1 | vasoactive intestinal peptide receptor 1 | 10.83 | 0.038 |
| GALR1 | galanin receptor 1 | 10.77 | 0.039 |
| TMEM132A | transmembrane protein 132A | 10.66 | 0.039 |
| GABRG2 | gamma-aminobutyric acid (GABA) A receptor, gamma 2 | 10.6 | 0.039 |
| ABHD8 | abhydrolase domain containing 8 | 10.47 | 0.040 |
| POU6F2 | POU class 6 homeobox 2 | 10.42 | 0.040 |
| PTPN3 | protein tyrosine phosphatase, non-receptor type 3 | 10.4 | 0.040 |
| VIPR2 | vasoactive intestinal peptide receptor 2 | 10.4 | 0.040 |
| KCNS2 | potassium voltage-gated channel, delayed-rectifier, | 10.38 | 0.040 |
| EPHX4 | epoxide hydrolase 4 | 10.35 | 0.040 |
| TCHH | trichohyalin | 10.29 | 0.040 |
| SDR16C5 | short chain dehydrogenase/reductase family 16C, member | 10.28 | 0.040 |
| MFSD3 | major facilitator superfamily domain containing 3 | 10.22 | 0.041 |
| DNM1 | dynamin 1 | 10.15 | 0.041 |
| RASIP1 | Ras interacting protein 1 | 10.08 | 0.042 |
| MCHR2 | melanin-concentrating hormone receptor 2 | 10.03 | 0.042 |
| SMPX | small muscle protein, X-linked | 9.95 | 0.043 |
| TNNI3K | TNNI3 interacting kinase | 9.94 | 0.043 |
| CIT | citron (rho-interacting, serine/threonine kinase 21) | 9.83 | 0.044 |
| CHRD | chordin | 9.78 | 0.044 |
| HRH2 | histamine receptor H2 | 9.76 | 0.044 |
| MGP | matrix Gla protein | 9.72 | 0.044 |
| CCDC64B | coiled-coil domain containing 64B | 9.71 | 0.044 |
| GRIN2A | glutamate receptor, ionotropic, N-methyl D-aspartate 2A | 9.62 | 0.044 |
| EPHB6 | EPH receptor B6 | 9.61 | 0.044 |
| NIPAL2 | NIPA-like domain containing 2 | 9.6 | 0.044 |
| CASQ1 | calsequestrin 1 (fast-twitch, skeletal muscle) | 9.56 | 0.044 |
| SHD | Src homology 2 domain containing transforming protein D | 9.54 | 0.044 |
| TNNC2 | troponin C type 2 (fast) | 9.51 | 0.044 |
| WFDC2 | WAP four-disulfide core domain 2 | 9.51 | 0.044 |
| ADCY1 | adenylate cyclase 1 (brain) | 9.47 | 0.044 |
| PAH | phenylalanine hydroxylase | 9.44 | 0.044 |
| LINGO2 | leucine rich repeat and Ig domain containing 2 | 9.44 | 0.044 |
| CCKBR | cholecystokinin B receptor | 9.42 | 0.044 |
| DNAJC5G | DnaJ (Hsp40) homolog, subfamily C, member 5 gamma | 9.4 | 0.044 |
| HECW1 | HECT, C2 and WW domain containing E3 ubiquitin protein ligase | 9.37 | 0.044 |
| EFNA5 | ephrin-A5 | 9.24 | 0.047 |
| GSTT2 | glutathione S-transferase theta 2 | 9.23 | 0.047 |
| PREP | prolyl endopeptidase | 9.15 | 0.048 |
| ELAVL4 | ELAV (embryonic lethal, abnormal vision, Drosophila)-like 4 (Hu | 9.07 | 0.048 |
| CPNE9 | copine family member IX | 9.04 | 0.048 |
| PRSS3 | protease, serine, 3 | 9.02 | 0.048 |
| LRTM2 | leucine-rich repeats and transmembrane domains 2 | 8.98 | 0.048 |
| RCAN2 | regulator of calcineurin 2 | 8.96 | 0.048 |
| IPCEF1 | interaction protein for cytohesin exchange factors 1 | 8.93 | 0.048 |
| CA4 | carbonic anhydrase IV | 8.86 | 0.048 |
| THEMIS | thymocyte selection associated | 8.86 | 0.048 |
| KCNH5 | potassium voltage-gated channel, subfamily H (eag-related), | 8.84 | 0.048 |
| RIMS3 | regulating synaptic membrane exocytosis 3 | 8.83 | 0.048 |
| SRPK1 | SRSF protein kinase 1 | 8.77 | 0.048 |
| FES | feline sarcoma oncogene | 8.77 | 0.048 |
| PAK1 | p21 protein (Cdc42/Rac)-activated kinase 1 | 8.76 | 0.048 |
| SPATS2L | spermatogenesis associated, serine-rich 2-like | 8.72 | 0.048 |
| FBN1 | fibrillin 1 | 8.71 | 0.048 |
| HTR1F | 5-hydroxytryptamine (serotonin) receptor 1F, G protein-coupled | 8.71 | 0.048 |
| ZSWIM3 | zinc finger, SWIM-type containing 3 | 8.69 | 0.048 |
| LYPD5 | LY6/PLAUR domain containing 5 | 8.68 | 0.048 |
| PHLDA2 | pleckstrin homology-like domain, family A, member 2 | 8.62 | 0.049 |
| NUAK1 | NUAK family, SNF1-like kinase, 1 | 8.62 | 0.049 |
| CITED4 | Cbp/p300-interacting transactivator, with Glu/Asp-rich carboxy- | 8.62 | 0.049 |
| ITPR1 | inositol 1,4,5-trisphosphate receptor, type 1 | 8.57 | 0.050 |
| ZDHHC22 | zinc finger, DHHC-type containing 22 | 8.52 | 0.050 |
